# Increased rates of genomic mutation in a biofilm co-culture model of *Pseudomonas aeruginosa* and *Staphylococcus aureus*

**DOI:** 10.1101/387233

**Authors:** C.J. Frapwell, R.P. Howlin, O. Soren, B.T. McDonagh, C.M. Duignan, R.N. Allan, A.R. Horswill, P. Stoodley, Y. Hu, A.R.M. Coates, J. S. Webb

## Abstract

Biofilms are major contributors to disease chronicity and are typically multi-species in nature. *Pseudomonas aeruginosa* and *Staphylococcus aureus* are leading causes of morbidity and mortality in a variety of chronic diseases but current *in vitro* dual-species biofilms models involving these pathogens are limited by short co-culture times (24 to 48 hours). Here, we describe the establishment of a stable (240 hour) co-culture biofilm model of *P. aeruginosa* and *S. aureus* that is reproducible and more representative of chronic disease.

The ability of two *P. aeruginosa* strains, (PAO1 and a cystic fibrosis isolate, PA21), to form co-culture biofilms with *S. aureus* was investigated. Co-culture was stable for longer periods using *P. aeruginosa* PA21 and *S. aureus* viability within the model improved in the presence of exogenous hemin. Biofilm co-culture was associated with increased tolerance of *P. aeruginosa* to tobramycin and increased susceptibility of *S. aureus* to tobramycin and a novel antimicrobial, HT61, previously shown to be more effective against non-dividing cultures of *Staphylococcal spp.* Biofilm growth was also associated with increased short-term mutation rates; 10-fold for *P. aeruginosa* and 500-fold for *S. aureus*.

By describing a reproducible 240 hour co-culture biofilm model of *P. aeruginosa* and *S. aureus*, we have shown that interspecies interactions between these organisms may influence short-term mutation rates and evolution, which could be of importance in understanding the adaptive processes that lead to the development of antimicrobial resistance.

## Introduction

Treatment of bacterial infections is often complicated by the presence of biofilms; communities of bacteria characterised by a heterogeneous composition and tolerance to antimicrobial treatment^1,2^. Tolerance towards antimicrobial compounds has further been linked with the emergence of mutations that confer AMR in planktonic cultures^3^ so it is possible that biofilm-mediated tolerance mechanisms could contribute similarly. Therefore, the development of relevant biofilm models is vital to understanding the interplay between biofilm tolerance mechanisms and the emergence of AMR.

Two bacterial species that are commonly implicated in biofilm infections are *Pseudomonas aeruginosa* and *Staphylococcus aureus.* In cystic fibrosis co-infection is associated with increased inflammation and reduced therapeutic outcomes for patients^4^, and in chronic wounds they are the most commonly co-isolated bacterial species and linked to poorer clinical outcomes^5^. However, whether the two species are co-localised or spatially partitioned remains a point of contention; in part because *in vitro* studies suggest that the relationship of these two bacteria is often antagonistic in nature^6–8^.

Although it is widely recognised that *in vivo* biofilms are often composed of a multispecies consortium, the majority of *in vitro* biofilm studies fail to reflect this, focusing on single species. Previous models investigating co-culture of *P. aeruginosa* and *S. aureus in vitro* have frequently observed that *P. aeruginosa* rapidly outcompetes and reduces *S. aureus* viability within 24 hours^7,8^. Consequently, use of these species within *in vitro* co-culture biofilm models is often restricted to short incubation periods, such as 24 or 48 hours^7–10^, which is not representative of long-term biofilm colonisation associated with chronic infection. Furthermore, use of these short-term *in vitro* models does not address or investigate factors that could improve the viability of *S. aureus* within a co-culture population.

There is an urgent need to investigate the impact of interspecies interactions within biofilms on bacterial persistence, virulence and evolvability in order to develop novel treatment strategies and circumvent the emergence of adaptive mechanisms, such as those associated with AMR. In this study we aimed to develop and characterise an *in vitro* dual-species biofilm formed by *S. aureus* and *P. aeruginosa* that is more representative of chronic infection. A 240-hour co-culture model was established and used to determine the impact on the antimicrobial susceptibility and individual mutation rates of both bacterial species. To our knowledge, this is the first documented approach using a fluctuation assay to assess short-term biofilm evolvability.

## Materials and Methods

### Bacterial Strains and Growth Conditions

The species/strains utilised in this study were *P. aeruginosa* PAO1, the cystic fibrosis isolate *P. aeruginosa* PA21, *S. aureus* UAMS-1 and *S. aureus* USA 300 LAC AH1279 (supplementary Table 1 for more information). Overnight planktonic cultures of *P. aeruginosa* and *S. aureus* were grown in Luria-Bertani broth, (LB, ForMedium, UK) and Tryptic Soy Broth, (TSB, Oxoid, UK), respectively. Biofilm cultures were grown in either Nunc Coated 6 well polystyrene plates (Thermo-Scientific, UK) for biomass experiments or poly-L-lysine coated glass bottomed dishes (MatTek, USA), for imaging experiments. Cultures were grown aerobically at 37 °C, with agitation at 120 rpm for planktonic cultures and 50 rpm for biofilms.

For enumeration and differentiation between *P. aeruginosa* and *S. aureus*, planktonic and biofilm cultures were plated onto either cetrimide agar (Oxoid, UK) supplemented with 1% glycerol (Sigma-Aldrich, UK) and Baird Parker agar (BPA), supplemented with 5% egg yolk tellurite emulsion (Oxoid, UK).

### Growth Kinetics

Overnight cultures were diluted to 10^6^ CFU ml^−1^ in LB or TSB as appropriate and incubated for 24 hours at 37 °C with OD_560_ measurements taken every 15 minutes for 15 hours using a 96 well plate reader (BMG Omega).

### Crystal Violet Assay

*P. aeruginosa* and *S. aureus* biofilms were grown for 72 hours using LB or TSB as appropriate, with fresh media exchanges every 24 hours. At 24, 48 and 72-hour time points spent media was removed and biofilms stained with 0.1% (v/v in dH_2_O) crystal violet for 10 minutes at room temperature. Biofilms were rinsed 3 times with dH_2_O then 30% acetic acid added to resolubilise the crystal violet. After 10-minute incubation at room temperature, with light shaking, the OD_550_ of the crystal violet suspension was measured using a spectrophotometer (Jenway 6300), with 30% acetic acid used as a blank.

### Planktonic Competition Assays

The relative fitness of both *S. aureus* strains was determined against both *P. aeruginosa* strains in 20 % BHI (Oxoid, UK) or 20 % BHI supplemented with hemin (Sigma-Aldrich, UK) at a final concentration of 2, 20 or 100 μM. Overnight cultures were diluted to 10^6^ CFU ml^−1^ in the appropriate medium to enable co-culture of the *P. aeruginosa* and *S. aureus* strains at a 1:1 ratio. Cultures were incubated at 37 °C, 120 rpm for 24 hours. Initial and endpoint cell number were obtained by serially diluting in Hanks Balanced Salt Solution (HBSS, Sigma-Aldrich UK) prior to plating on cetrimide agar and BPA. Plates were incubated at 37 °C for 24 hours.

The relative fitness of each bacterial species and strain was obtained by comparing the ratio of their Malthusian parameters (M_*P.* aeruginosa_ and M_*S.* aureus_) whereby;

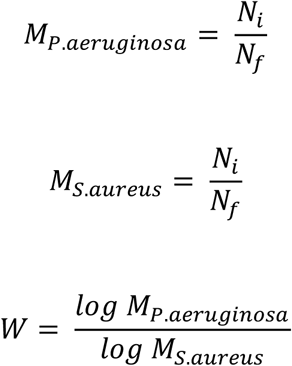

*N*_*i*_ = Initial cell number

*N*_*f*_ = Final cell number

*W =* Relative fitness

### Biofilm Co-Culture Optimisation

The ability for *P. aeruginosa* and *S. aureus* to form dual species biofilms was assessed over 240 hours by adapting a previous method^11^. Overnight cultures of *P. aeruginosa* and *S. aureus* were diluted to 10^5^ CFU ml^−1^ in ½ strength BHI at a 1:1 ratio of each species. 1 ml was used to inoculate each well of a Nunclon coated 6 well plate (Thermo Scientific, UK) and incubated for 6 hours at 37 °C, 50 rpm to facilitate bacterial attachment. Media was then replaced with 4 ml of 20% BHI or 20% BHI supplemented with hemin (Sigma-Aldrich, UK) at a final concentration of 2, 20 or 100 μM. Media was replaced after a further 18 hours, then every 24 hours thereafter. After 24, 72, 168 and 240 hours, biofilms were rinsed twice with HBSS to remove non-adherent cells and harvested using a cell scraper. Cell suspensions were serially diluted in HBSS and plated onto cetrimide agar and BPA for selective enumeration of *P. aeruginosa* and *S. aureus*, respectively.

### Confocal Laser Scanning Microscopy of *P. aeruginosa* and *S. aureus* biofilms

Mono- and co-culture biofilms of *P. aeruginosa* PA21 with *S. aureus* UAMS-1 or *S. aureus* LAC were cultured to assess biofilm architecture. Biofilms were grown using 20% BHI supplemented with 20 μM hemin in MatTek dishes. Biofilms were assessed at 24, 72, 168 and 240 hours of growth. Prior to imaging, spent media was removed and the biofilms rinsed twice with HBSS before staining for 15 minutes with 1 ml of LIVE *Bac*Light Bacterial Gram stain (Life Technologies), (3 μl ml^−1^ SYTO9, 2 μl ml^−1^ hexidium iodide). Imaging was performed using an inverted Leica TCS SP8 confocal laser scanning microscope and a 63x glycerol immersion lens, with 1 μm vertical sections. Fluorescent dyes were excited using concurrent 514 nm and 561 nm lasers.

### Antimicrobial Susceptibility Testing

The minimum inhibitory concentration (MIC) of rifampicin (Sigma-Aldrich, UK) was determined for *P. aeruginosa* PA21 and *S. aureus* UAMS-1 using the broth microdilution method^12^, with a two-fold dilution series of rifampicin (0 to 128 μg ml^−1^). Following a 24-hour incubation at 37° C, the endpoint OD_680_ was measured with a microplate reader (BMG Omega). The MIC was the antimicrobial concentration that resulted in no bacterial growth.

The biofilm minimum bactericidal concentrations (MBC) of tobramycin, vancomycin and HT61^12^, was determined for mono- and co-culture biofilms of *P. aeruginosa* PA21 and *S. aureus* UAMS-1 using a method adapted from Howlin *et al* (2015)^13^. Biofilms were cultured for 72 hours in Nunclon coated 6 well plates as previously described using 20% BHI supplemented with 20 μM hemin. Spent media was replaced with antimicrobial supplemented media (two-fold dilution series between 0 and 128 μg ml^−1^). After an additional 24 hours of incubation, biofilms were rinsed twice with HBSS, harvested with a cell scraper and serially diluted and plated onto cetrimide agar and BPA. The biofilm MBCs were identified as the concentration leading to a 3-log reduction in CFU’s.

### Estimation of Planktonic and Biofilm Mutation Rates

Fluctuation tests were performed for planktonic cultures as previously described in Foster (2006)^14^ and adapted for use with biofilm cultures grown in 6 well plates. Overnight cultures of *P. aeruginosa* PA21 and *S. aureus* UAMS-1 were diluted to 10^3^ CFU ml^−1^ in 50% BHI, either in isolation, or in a 1:1 co-culture. 30 parallel planktonic or biofilm cultures were initiated using 1 ml of the inoculum in 20 ml universal containers or Nunclon coated 6 well plates, respectively. All cultures were incubated at 37 °C, 50 rpm for 6 hours, with planktonic cultures tilted at approximately 45° to allow for media movement.

Planktonic cultures were centrifuged at 4000 × *g* for 15 minutes and the cell pellet resuspended in 4 ml of 20% BHI supplemented with 20 μM hemin. For biofilm cultures, spent media was replaced with 4 ml of 20% BHI supplemented with 20 μM hemin. Cultures were incubated at 37 °C, 50 rpm for a further 18 hours. Following incubation, planktonic cultures were centrifuged at 4000 x *g*, the cell pellet rinsed twice with HBSS, then re-suspended in 500 μl HBSS. Biofilm cultures were rinsed twice with HBSS, harvested using a cell scraper and resuspended into 500 μl HBSS.

Final cell counts were determined by plating 5 random cultures onto cetrimide agar and BPA. Half of each remaining planktonic and biofilm suspension was then plated onto cetrimide agar and BPA supplemented with 64 μg ml^−1^ or 0.25 μg ml^−1^ rifampicin (4 × calculated MIC for each species) for selection of spontaneous *P. aeruginosa* or *S. aureus* mutants, respectively. Plates were incubated at 37 °C and enumerated after 24 hours (plates without antibiotics) or 48 hours (rifampicin plates).

Mutation rates were calculated using FALCOR and the Ma-Sandri-Sarkar Maximum Likelihood Estimator^15^. Fluctuation tests were performed in biological duplicate (60 technical replicates).

### Genomic Comparison of *P. aeruginosa* PAO1 and PA21

Short read sequencing of both *P. aeruginosa* strains was performed by MicrobesNG on Illumina platforms using 250 bp paired end reads. Long read sequencing was performed using the MinIon sequencing platform (Oxford Nanopore, UK and the rapid barcoding kit as per manufacturer’s instructions. Long read data was basecalled using albacore and trimmed and demultiplexed using PoreChop (https://github.com/rrwick/Porechop) with default settings.

Hybrid assemblies were performed *de novo* using Unicycler^16^ in normal mode resulting and annotated using PROKKA^17^. Annotations were preserved and genomes compared using RAST^18,19^.

### Statistical Analysis

Statistical analyses were performed using GraphPad Prism version 7.0d for Mac. Crystal violet data and comparisons between single species biofilms were made using multiple t-tests with the Holm-Sidak correction. Effects of media composition and bacterial competition on fitness were analysed using a 2-way ANOVA. Kruskal-Wallis tests with Dunn’s multiple comparisons were used to analyse within time point comparisons of biofilm co-culture and for comparison of biofilm maximum thickness, derived from microscopy data. For all of the above statistical tests, α ≤ 0.05.

The R package, RSalvador^20^, was used to calculate 94% confidence intervals for the fluctuation test data. Confocal image z-stacks were analysed using the COMSTAT 2 plug in for ImageJ (downloadable at www.comstat.dk)^21^.

Statistical significance was determined between fluctuation tests by manually comparing 94 % confidence intervals (shown to mimic statistical tests for *p* ≤ 0.01 and a valid statistical approach when comparing fluctuation test data with differing terminal cell population sizes^22,23^.

## Results

### Planktonic co-culture of *P. aeruginosa* and *S. aureus* did not affect relative fitness of either species, despite differences in growth kinetics or biofilm formation

Basic phenotyping of each strain of *P. aeruginosa* and *S. aureus* was performed, comparing growth kinetics, (Figure 1A), capacity to form biofilms (Figure 1B), and the relative fitness of each strain when grown as planktonic co-cultures in a selection of defined media (Figure S1). No difference in growth kinetics was observed for either strain of *S. aureus*, and both exhibited similar biofilm-forming capacities with no statistically significant differences at any time point (multiple t-tests, Holm-Sidak correction; 24 hr *p* = 0.505, 48 hr *p* = 0.615, 72 hr *p* = 0.893).

**Figure 1:**
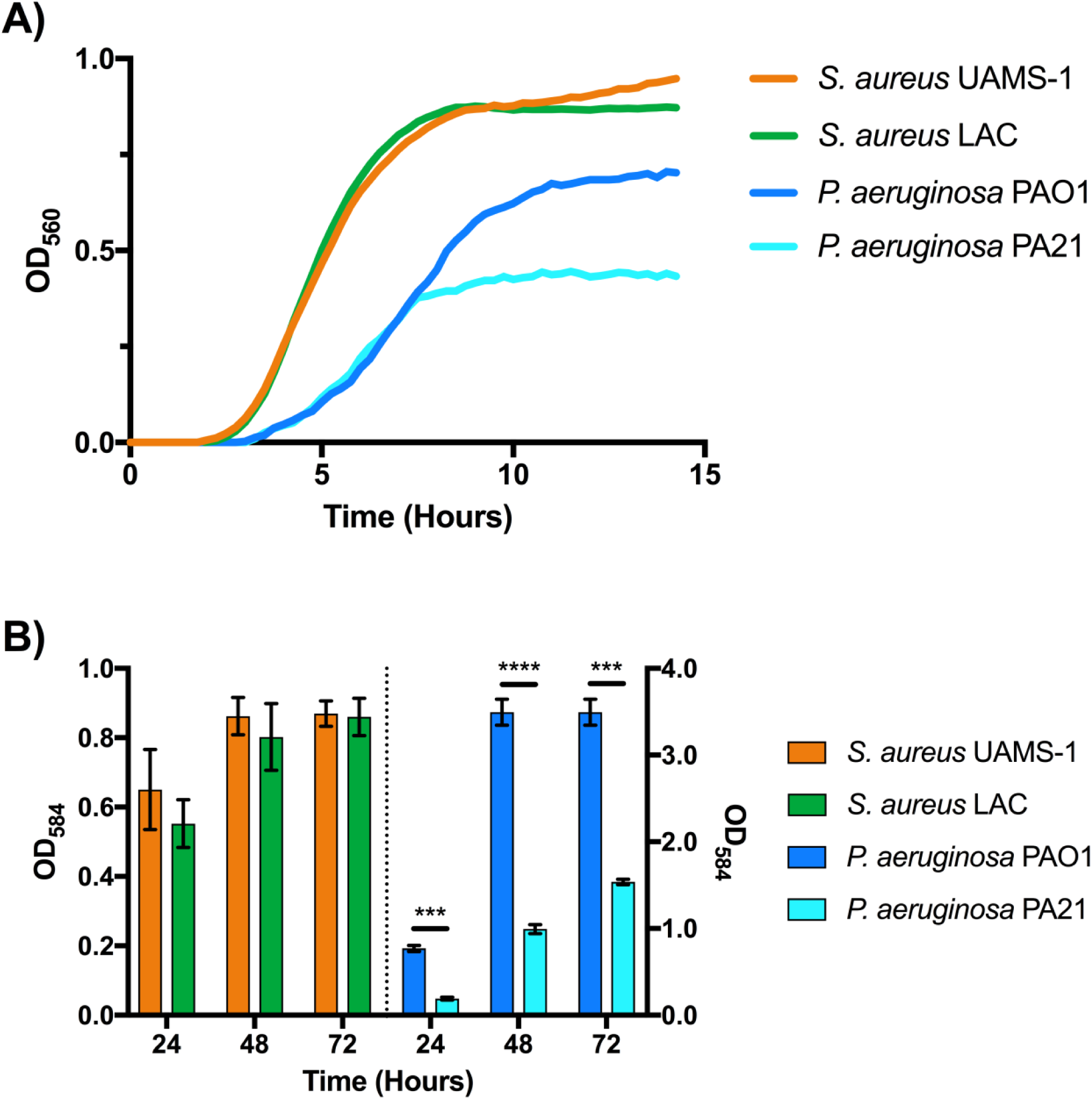
Growth Kinetics and Biofilm-forming Ability of P. aeruginosa PAO1, PA21 and S. aureus UAMS-1 and LAC. A) S. aureus. UAMS-1 and LAC exhibit near identical growth kinetics and biofilm formation. *P. aeruginosa* PAO1 and PA21 both enter exponential phase at the same point in time, but PA21 enters stationary earlier resulting in reduced culture density. B) Biofilm formation of PA21 is less than that measured for PAO1 at all time points. Growth curves: n = 12 for *S. aureus*, n = 3 for *P. aeruginosa*, Crystal violet assays: n = 3. Error bars represent standard error of the mean. **** *p* > 0.0001, *** *p* > 0.001. P values calculated using multiple unpaired student t-tests, corrected using the Holm-Sidak method.

For *P. aeruginosa*, the exponential phase of growth for PAO1 was approximately 2 hours longer than that of PA21, although final cell density was equal between cultures (data not shown). PAO1 formed biofilms with considerably more biomass at each measured time point compared to PA21, (multiple t-tests, Holm-Sidak correction; 24 hr *p* = 0.0001, 48 hr *p* < 0.0001, 72 hr *p* = 0.0002). Both strains formed more biofilm biomass than either strain of *S. aureus*.

Iron availability is known to be important for the growth of pathogens within the cystic fibrosis lung^24,25^ and hemin was chosen to mimic a potential *in vivo* source of ferric iron donor. Planktonic relative fitness was determined in 20% BHI supplemented with hemin to a final concentration of either 2, 20 or 100 μM. Relative fitness was equal to 1 for each combination in all media compositions with no statistical differences between any calculated values, suggesting no decrease in fitness for either species during planktonic co-culture (2-way ANOVA, interaction *p* = 0.0538, bacterial combinations *p =* 0.1960, media choice *p =* 0.5100)

### Addition of Hemin Does Not Affect Viability of *S. aureus* and *P. aeruginosa* During Growth as a Single Species Biofilm

Single species biofilms of each *P. aeruginosa* strain were unaffected by supplementation of media with hemin, although there was intra-strain variation (see supplementary information, figure S2). *P. aeruginosa* PA21 formed biofilms with a lower cell density compared to those formed by *P. aeruginosa* PAO1 at 24 hours in all media tested (multiple unpaired t-tests at for each media combination, *p* < 0.05). However, by 240 hours of growth any difference in cell density was statistically insignificant (*p >* 0.05)

Both *S. aureus* UAMS-1 and *S. aureus* LAC formed biofilms with equal cell densities at 24 and 72 hours, regardless of the hemin concentration in the media. However, by 168 hours, regardless of hemin supplementation, *S. aureus* LAC biofilms were approximately 1 log lower cell density compared to *S. aureus* UAMS-1, a decrease that was still apparent following 240 hours of growth (*p* < 0.05).

### *P. aeruginosa* strain selection and increased hemin concentrations can improve *S. aureus* survival during Biofilm Co-Culture

Co-culture biofilms of *P. aeruginosa* PAO1 or PA21 with *S. aureus* UAMS-1 or LAC were grown for 10 days (240 hours) and the impact of different hemin concentrations on the viability of the co-cultures assessed (Figure 2). *P. aeruginosa* viability was not affected by the addition of hemin to the media. While hemin improved *S. aureus* viability, it was present at lower abundance than *P. aeruginosa* in all co-cultures.

**Figure 2:**
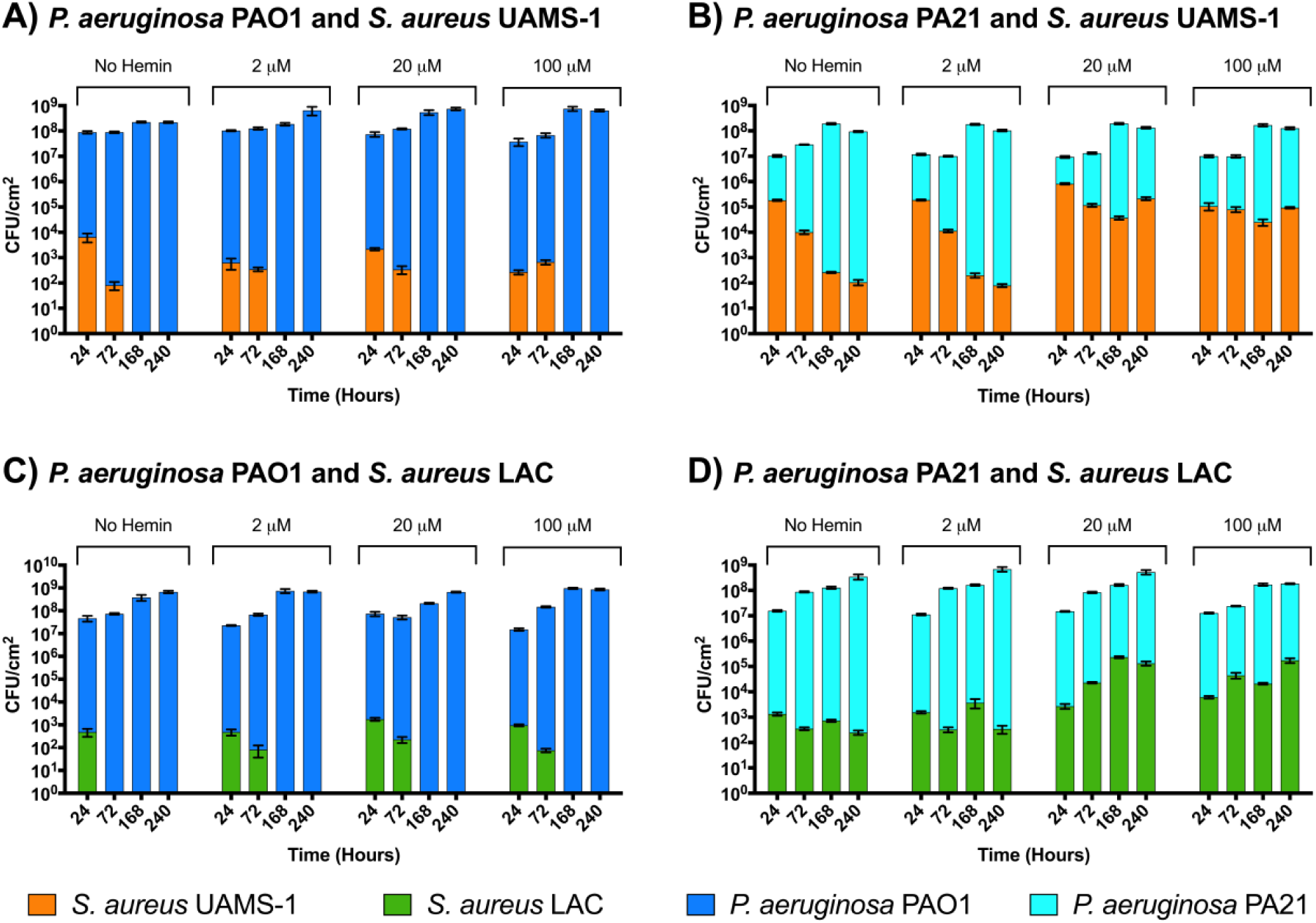
Effects of hemin supplementation on the dynamics of P. aeruginosa and S. aureus biofilm co-culture. The ability of S. aureus to survive in co-culture depends on a combination of the P. aeruginosa strain and the hemin concentration of the media. Both strains of S. aureus were less successful in co-culture with P. aeruginosa PAO1, compared to P. aeruginosa PA21. Addition of hemin at 2 μM facilitated the growth of S. aureus LAC with P. aeruginosa PAO1 for at least 72 hours and addition of hemin at either 20 μM or 100 μM significantly improved the viability of both S. aureus UAMS-1 and S. aureus LAC when in co-culture with P. aeruginosa PA21. Kruskal Wallis with Dunn’s multiple comparison performed to determine significance between different media compositions. Points of note are mentioned in the text. Due to the large number of comparisons, significance levels are not presented on chart and are instead compiled in the supplementary information. *n = 9 from 3 biological replicates*

When *S. aureus* UAMS-1 was co-cultured with *P. aeruginosa* PAO1, no viable *S. aureus* cells were identified after 168 hours of growth, regardless of hemin supplementation. Hemin supplementation slightly reduced UAMS-1 viability at 24 hours, although this decrease was not statistically significant with 20 μM hemin (Figure 2, panel A, 24 hour *p* values; No hemin vs 2 μM = 0.0167, No hemin vs 20 μM > 0.999, No hemin vs 100 μM = 0.0006).

*S. aureus* LAC was not detectable following 72 hours of growth with *P. aeruginosa* PAO1. Hemin supplementation improved *S. aureus* viability so that it was detectable at 72 hours, although due to a number of zero counts in 2 μM hemin, the improved viability was only significant in 20 or 100 μM hemin (72 hour *p* values No hemin vs 2 μM = 0.7089, No hemin vs 20 μM = 0.0015, No hemin vs 100 μM = 0.0241). However, similar to *S. aureus* UAMS-1, *S. aureus* LAC was not detectable after 168 hours regardless of hemin supplementation.

Conversely, both strains of *S. aureus* were detectable after 240 hours of co-culture with *P. aeruginosa* PA21 (Figure 2, panels B and D). Hemin supplementation improved viability further, although for both strains, *S. aureus* counts in 2 μM hemin were statistically identical to counts in non-supplemented media at all time points (*p* > 0.9999). In the non-supplemented and 2 μM hemin supplemented media *S. aureus* UAMS-1 viability decreased over 240 hours from approx. 10^5^ CFU cm^−2^ to 10^2^ CFU cm^−2^, whereas *S. aureus* LAC remained at a density between approximately 10^2^ and 10^3^ CFU cm^−2^. Supplementation of media with 20 and 100 μM hemin increased *S. aureus* viability compared to non-supplemented media with final counts at 240 hours approximately 10^5^ CFU cm^−2^ for both strains (*S. aureus* No hemin vs 20 μM *p* values; UAMS-1 240 hour < 0.0001, LAC 240 hour = 0.0033). *S. aureus* viability in media supplemented with 100 μM hemin was similar to that in media supplemented with 20 μM hemin, with the exception of *S. aureus* UAMS-1 at 24 hours, which was 1 log lower in density (*p* < 0.0001). All other differences were statistically insignificant (20 μM vs 100 μM *S. aureus* UAMS-1 *p* values; 72 hour > 0.9999, 168 hour > 0.9999, 240 hour = 0.5851 and 20 μM vs 100 μM *S. aureus* LAC *p* values; 24 hour = 0.0753, 72 hour > 0.9999, 168 hour = 0.4190, 240 hour > 0.9999)

Addition of 100 μM hemin did not discernibly improve viability of *S. aureus* or *P. aeruginosa* compared to supplementation with 20 μM hemin. For this reason, subsequent assays utilised hemin at a concentration of 20 μM in 20 % BHI.

### Visualisation of *P. aeruginosa* and *S. aureus* Biofilm Co-Cultures

Representative images are presented in Figure 3 (*P. aeruginosa* PA21 and *S. aureus* UAMS-1) and Figure 4, (*P. aeruginosa* PA21 and *S. aureus* LAC).

**Figure 3:**
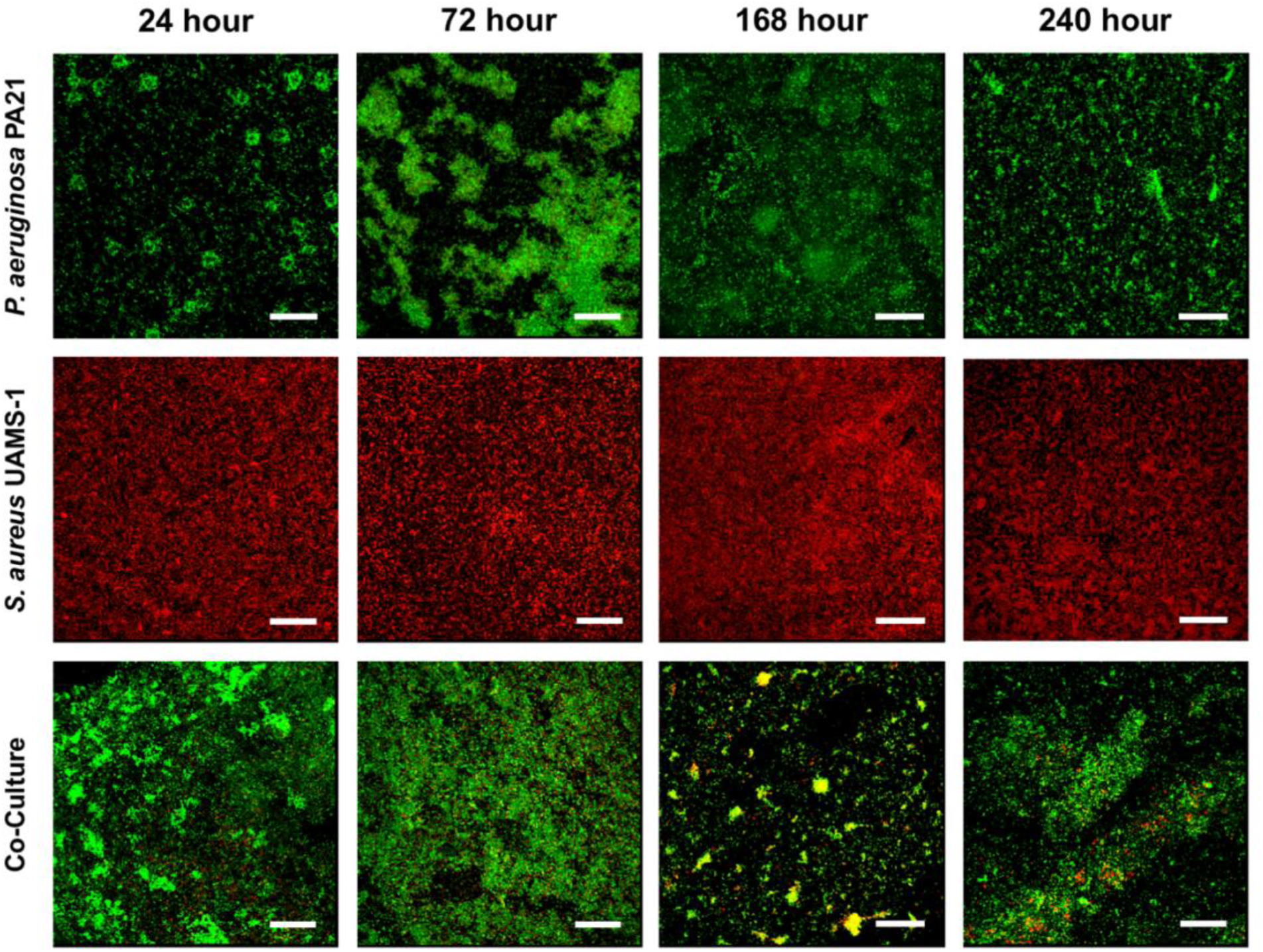
Representative Confocal Laser Scanning Microscopy images of a single and dual species biofilms of P. aeruginosa PA21 and S. aureus UAMS-1 over 240 hours. Biofilms were stained with a fluorescent Gram stain for differentiation between species. Hexidium iodide stains Gram positive bacteria to fluoresce red, (S. aureus), while SYTO9 counterstains the remaining Gram negative bacteria (P. aeruginosa) and fluoresces green. Scale bars represent 25 μm.

**Figure 4:**
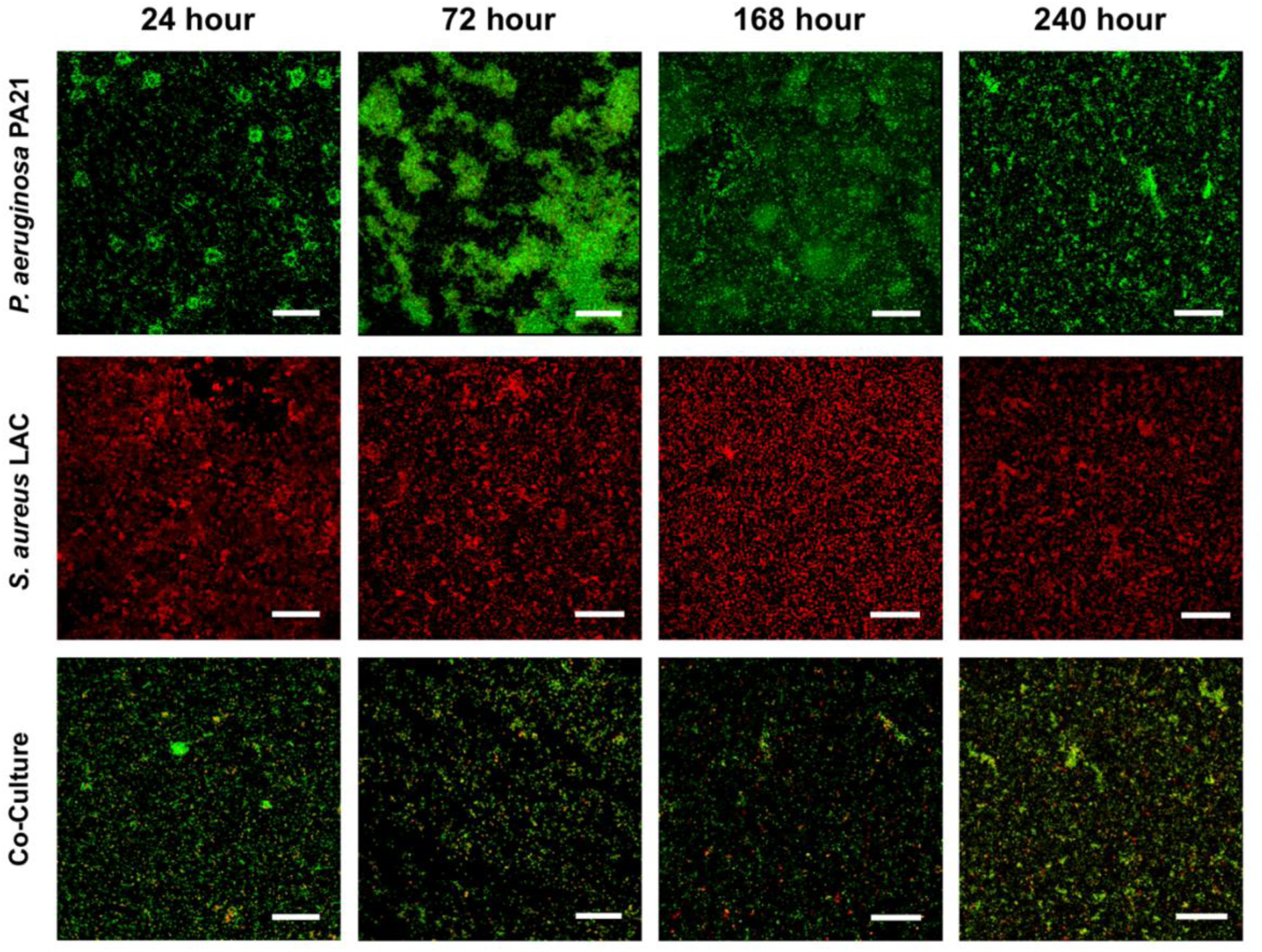
Representative Confocal Laser Scanning Microscopy images of a single and dual species biofilms of P. aeruginosa PA21 and S. aureus LAC over 240 hours. Biofilms were stained with a fluorescent Gram stain for differentiation between species. Hexidium iodide stains Gram positive bacteria to fluoresce red, (S. aureus), while SYTO9 counterstains the remaining Gram-negative bacteria (P. aeruginosa). Scale bars represent 25 μm.

When cultured in isolation, *P. aeruginosa* PA21 forms microcolony structures after 24 hours of growth. These structures expand after 72 hours and by 168 hours of growth have formed a confluent layer. By 240 hours the overall density of the biofilm appears to have reduced, although the remaining microcolonies do contribute to increased biofilm thickness (see supplementary information, Figure S3). Both *S. aureus* UAMS-1 and *S. aureus* LAC undergo similar biofilm development, corroborating the previously obtained crystal violet and CFU data (Figure 1 and Figure S2, respectively), forming a uniform sheet of biomass without any significant changes in maximum biofilm thickness over 240 hours (Figure S3, *p* > 0.05).

During co-culture of *P. aeruginosa* PA21 and *S. aureus* UAMS-1, *S. aureus* is distributed uniformly throughout the biofilm for the first 72 hours. By 168 hours of growth, *S. aureus* appears to have clustered around cellular aggregates, (indicated by the yellow colour within the image resulting from a high number of green and red cells in close proximity, similar to the observation made by DeLeon *et al*^10^. By 240 hours, the dense areas of *Staphylococcal* cells appear to have lessened, although it still found throughout the biofilm. Co-culture of *S. aureus* LAC with *P. aeruginosa* PA21 shows a similar morphology over the first 72 hours of growth. Unlike *S. aureus* UAMS-1, *S. aureus* LAC remains uniformly distributed at 168 and 240 hours with far fewer localised aggregates.

### Biofilm Co-Culture Alters the Antimicrobial Susceptibility of *P. aeruginosa* and *S. aureus*

**Table 1:** Antimicrobial susceptibility of P. aeruginosa PA21 and S. aureus UAMS-1 in single and dual species biofilms. Biofilm minimum bactericidal concentrations (MBCs) of tobramycin and vancomycin defined as the concentration of antimicrobial that reduced the viable counts of each species by 3 log or more. Values in table represent antimicrobial concentration in μg ml^−1^. Biofilm co-culture increased the tobramycin MBC for P. aeruginosa from 2 μg ml^−1^ to 4 μg ml^−1^ and decreased the Biofilm MBC of S. aureus from 16 μg ml^−1^ to 2 μg ml^−1^. HT61 was not effective against P. aeruginosa, however biofilm co-culture increased S. aureus susceptibility four-fold (Single species MBC = 64 μg ml^−1^ Co-culture MBC = 16 μg ml^−1^) Co-culture did not alter susceptibility of either species to vancomycin. n = 9, from 3 biological replicates.

| Antimicrobial | *P. aeruginosa* PA21 |  | *S. aureus* UAMS-1 |  |
| --- | --- | --- | --- | --- |
|  | Single Species | Dual Species | Single Species | Dual Species |
| Tobramycin | 2 | 4 | 16 | 2 |
| Vancomycin | > 128 | > 128 | > 128 | > 128 |
| HT61 | > 128 | > 128 | 64 | 16 |

To demonstrate that the biofilm model could be utilised in phenotyping experiments the antimicrobial susceptibility of established mono- and co-culture biofilms of *S. aureus* UAMS-1 and *P. aeruginosa* PA21 was determined using two antimicrobials in clinical use (tobramycin and vancomycin) and a novel antimicrobial compound currently in development (HT61).

Whereas vancomycin and HT61 had no effect on *P. aeruginosa* viability when grown as either a single or dual-species biofilm, the MBC of tobramycin increased from 2 to 4 μg ml^−1^. Similarly, *S. aureus* viability was not affected by vancomycin in either single or dual-species biofilms, however, the tobramycin and HT61 MBCs were reduced eightfold (16 to 2 μg ml^−1^) and fourfold (64 to 16 μg ml^−1^) respectively.

### Biofilm co-culture of *P. aeruginosa* and *S. aureus* significantly increases the mutation rate of each species

Understanding whether interspecies interactions can alter bacterial evolvability is incredibly relevant, considering the rapid emergence of AMR. The Luria-Delbrück fluctuation test was applied to both single species and dual species co-cultures of *P. aeruginosa* PA21 and *S. aureus* UAMS-1, both in planktonic and biofilm culture to measure the spontaneous mutation rate (via the development of rifampicin resistance) of each species (Figure 5).

**Figure 5:**
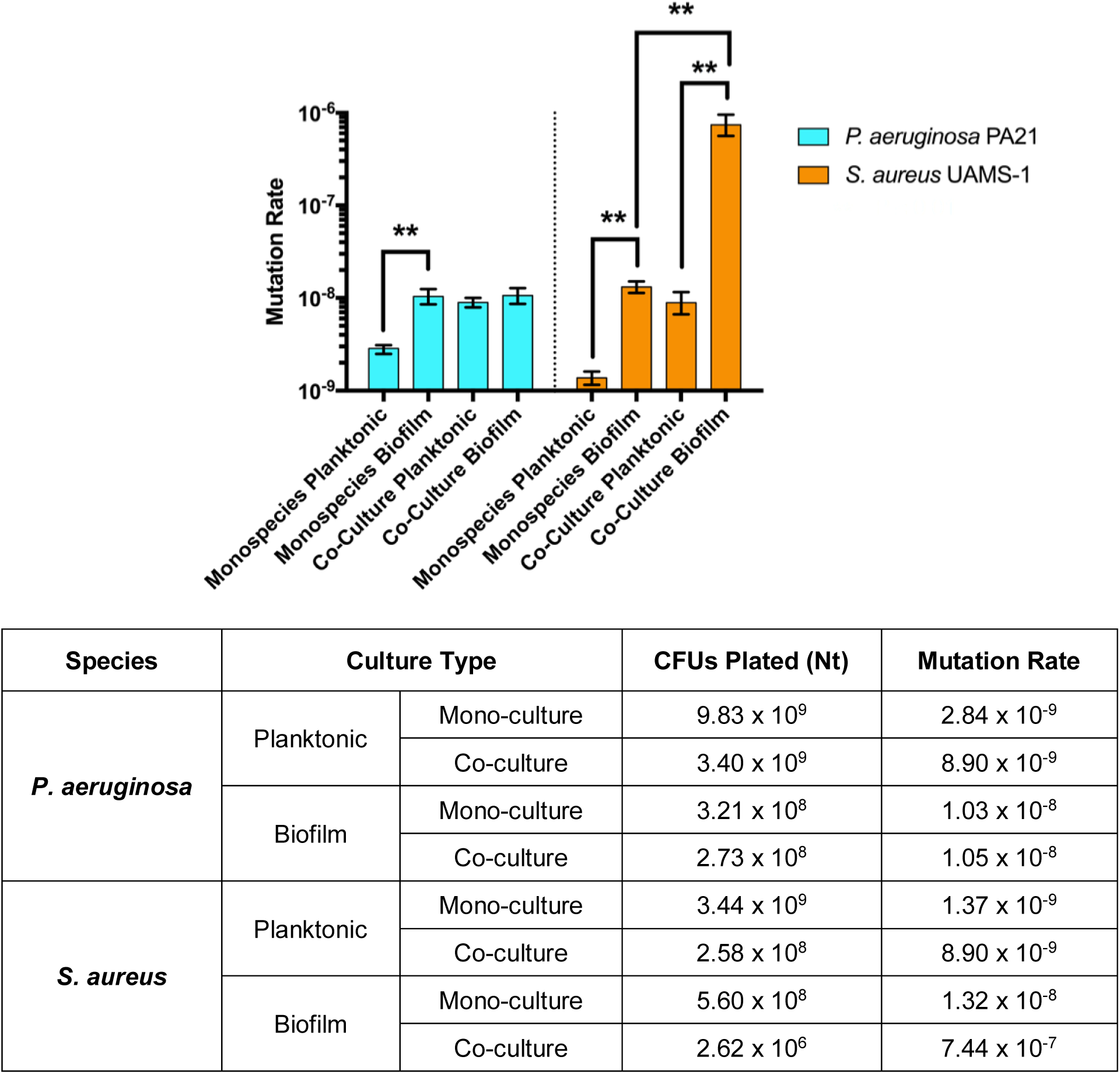
Effect of Biofilm Growth on Mutation Rate of P. aeruginosa PA21 and S. aureus UAMS-1. Following biofilm mono-culture, both species are shown to have an increased rate of mutation, (P. aeruginosa 1.03 × 10^−8^, S. aureus 1.32 × 10^−8^). The rate of mutation for P. aeruginosa stays at this level following co-culture planktonic growth and co-culture biofilm growth suggesting that while species interactions are important, in this case, the species interactions between P. aeruginosa and S. aureus do not impact the evolvability of P. aeruginosa more than the conditions of biofilm growth. The mutation rate of S. aureus following planktonic co-culture is approximately the same as that following biofilm mono-culture (Planktonic co-culture: 8.90 × 10^−9^, Biofilm mono-culture: 1.32 × 10^−8^), suggesting that the pressures associated with the presence of planktonic P. aeruginosa on staphylococcal evolvability is similar to those associated with mono-culture biofilm growth. However, following biofilm co-culture, the mutation rate of S. aureus is highly elevated to 7.44 × 10^−7^. This suggests that both biofilm growth and the interspecies interactions present following biofilm co-culture with P. aeruginosa may be important to understanding the evolvability of S. aureus. Error bars represent 94% confidence intervals. ** represents p ≤ 0.01, obtained by comparing overlap of confidence intervals. This method is a valid comparison as calculation of confidence intervals and accounts for differences in the terminal population density^23^

In planktonic mono-culture, the mutation rates of *P. aeruginosa* PA21 and *S. aureus* UAMS-1 were low, at 2.84 × 10^−9^ and 1.37 10^−9^ mutations per cell division, respectively. Planktonic co-culture led to an increase in mutation rate for both species to 8.90 × 10^−9^ mutations per cell division. Biofilm mono-culture resulted in a similar increase to 1.03 × 10^−8^ and 1.32 × 10^−8^ mutations per cell division for *P. aeruginosa* PA21 and *S. aureus* UAMS-1, respectively. Following biofilm co-culture, the mutation rate of *P. aeruginosa* remained at a similar level (1.05 × 10^−8^ mutations per cell division). However, the mutation rate of *S. aureus* increased to 7.44 × 10^−7^ mutations per cell division, which is an approximately 500-fold relative increase compared to the rate during planktonic mono-culture.

### *P. aeruginosa* Genome Comparison Reveals Strain Specific Features

Due to the differing abilities of *P. aeruginosa* PAO1 and *P. aeruginosa* PA21 to form co-culture biofilms, the genomes of these two strains were sequenced and compared to identify whether any obvious genetic differences might explain the different co-culture phenotypes. Assembly statistics and a complete list of differential genomic features are presented in the supplementary information.

## Genes Absent in *P. aeruginosa* PA21

77 genes were identified that were present only in *P. aeruginosa* PAO1. Of these, 41 were hypothetical, resulting in 36 annotated genes (complete list in S1), which include numerous phage proteins, helicases, manganese catalase and a number of genes associated with lipopolysaccharide production such as the virulence associated Wzx and Wzy flippases.

## Additional Genes in *P. aeruginosa* PA21

500 genes were found exclusively in *P. aeruginosa* PA21. Of these, 234 coded for hypothetical proteins while 266 genes were annotated across multiple categories including virulence factors, resistance genes, cell signalling, metabolism, as well as extensive phage associated proteins and proteins associated with DNA recombination (complete list in S2). While there are no specific genes that appear to be directly associated with the improved ability of *P. aeruginosa* PA21 to form a stable co-culture biofilm with *S. aureus*, there are numerous proteins of interest such the phd-doc toxin antitoxin (TA) system as well as additional proteins associated with iron uptake, such as periplasmic TonB, important for siderophore transport and uptake^26^.

## Discussion and Conclusion

In this paper, we developed a long-term *in vitro* biofilm co-culture of *P. aeruginosa* and *S. aureus* that could be maintained for at least 10 days and is more representative of chronic infection compared to current 24-48 hour models. Following optimisation of the biofilm co-culture we determined how biofilm co-culture altered the evolvability of each bacterial species, and using a genomic based approach compared and identified features of *P. aeruginosa* that could be implicated in sustaining a biofilm co-culture with *S. aureus*.

When comparing planktonic and biofilm co-culture, we found that the viability of *S. aureus* was only negatively affected during biofilm growth. This was interesting as numerous studies have shown that the viability of *S. aureus* is negatively affected by *P. aeruginosa* even in planktonic culture^10,27–29^. However, Miller *et al* (2017) found that use of a more nutrient rich medium improved *S. aureus* survival^30^. This suggests, consistent with our results that media composition is an important consideration for the development of a co-culture model.

We found that both *P. aeruginosa* strain and exogenous hemin concentration impacted *S. aureus* survival during biofilm co-culture. Altered levels of *S. aureus* killing by *P. aeruginosa* has been linked to latter’s ability to form biofilms; those that form less biofilm are less prone to *S. aureus* killing^7^. Based on crystal violet staining, *P. aeruginosa* PAO1 formed more robust single species biofilms than *P. aeruginosa* PA21 and also reduced the number of viable *S. aureus* during biofilm co-culture. It has been demonstrated that *P. aeruginosa* isolates taken from patients co-infected with *P. aeruginosa* and *S. aureus* are less competitive towards *S. aureus*^8,27^. While the exact clinical background of *P. aeruginosa* PA21 in relation to *S. aureus* co-culture is not available, this could be a factor that favours its co-culture with *S. aureus*.

We showed that increasing the concentration of hemin improved *S. aureus* viability during co-culture with *P. aeruginosa* PA21. During biofilm co-culture, *S. aureus* is lysed and used as an iron source for *P. aeruginosa* in a *Pseudomonas* quinolone signal, PQS, mediated process that is decreased in iron rich environments^7,31^. By increasing the exogenous iron concentration, it is possible that *P. aeruginosa* PA21 PQS expression was reduced, reducing *S. aureus* lysis. Measuring the changes in expression of associated genes in *P. aeruginosa* PA21 such as *pqsA* and *pqsH*, with differing concentrations of hemin may provide insight into this scenario.

Sequencing of the two *P. aeruginosa* genomes identified features that could be important targets for further investigation. The presence of additional iron uptake components and lack of manganese catalase enzymes, (typical for bacteria occupying low iron environments/possess ineffective iron uptake mechanisms)^32^, may mean that PA21 at the uptake of exogenous iron than PAO1. If uptake is more efficient, that could negatively regulate the production of molecules that are produced to lyse *S. aureus* and utilise it as an iron source instead^7^. As an aside, it was found that *P. aeruginosa* PA21 still contained genes encoding the siderophores pyoverdine and pyochelin, *pvdA* and *pchE*, respectively. The presence of these genes has previously been associated with increased killing of *S. aureus* in co-culture^9^. As *P. aeruginosa* PA21 was less lethal to *S. aureus* than *P. aeruginosa* PAO1, it suggests these genes are not implicated and the improved survival is a result of a different mechanism. Expression of the phd-doc TA system has been linked with translation inhibition^33^. If this module is activated by PA21 during co-culture, it could facilitate *S. aureus* survival by slowing *P. aeruginosa* growth and/or increase *S. aureus* tolerance. Further investigation into these elements would be required.

Mutation frequency is known to be elevated in biofilms^34,35^, which is a measure of the abundance of mutants within a population. However, mutation frequency can be distorted by the expansion of lineages harbouring low probability, “jackpot mutations” that occur during the early stages of growth. On the other hand, mutation rate, which measures the number of mutations sustained by a cell during its lifetime, accounts for jackpot mutations and is overall, a more robust measurement^14^. By applying the fluctuation test to planktonic and biofilm mono- and co-cultures, we provide additional evidence of this. We show that interspecies interactions can modulate rates further and highlight the importance of understanding interspecies interactions within bacterial communities. It has been demonstrated that planktonic cultures of *P. aeruginosa* undergo a different evolutionary trajectory when cultured in the presence of *S. aureus*, obtaining mutations in lipopolysaccharide biosynthesis genes and increased resistance to β-lactam antimicrobials^36^. As such, understanding the impact of these complex community interactions could prove critical in limiting AMR.

Biofilm co-culture of *P. aeruginosa* and *S. aureus* caused the two species to present with different levels of antimicrobial susceptibility compared to growth as a single species biofilm. For this study, we chose to test the efficacy of tobramycin, an aminoglycoside that is effective against both *P. aeruginosa* and *S. aureus*, and vancomycin, an important glycopeptide utilised in the control of *Staphylococcal* infections. The novel antimicrobial HT61 which has shown activity against *Staphylococcus spp.* was also tested^12^.

Biofilm co-culture caused *P. aeruginosa* to become less susceptible to tobramycin and *S. aureus* to become more susceptible. These effects have both been previously documented. Interactions between *P. aeruginosa* derived Psl polysaccharide and *S. aureus* derived Staphylococcal protein A can cause aggregates of *P. aeruginosa* to form, decreasing overall susceptibility to tobramycin^37^. Conversely, *P. aeruginosa* rhamnolipids production can potentiate tobramycin uptake in *S. aureus* cells^38^. *P. aeruginosa* production of the endopeptidase LasA has been linked to increased vancomycin susceptibility in *S. aureus*^38^, which was not observed in this study.

*S. aureus* susceptibility to HT61 was also increased during biofilm co-culture. HT61 is more effective against stationary phase cells due to the introduction of anionic membrane components^12,39^. Mechanisms that decrease *S. aureus* growth rates could potentiate HT61. Vancomycin tolerance has been associated with a similar mechanism in *P. aeruginosa* and *S. aureus* co-cultures^40^.

### Limitations and Conclusions

The *in vitro* model described here is reproducible, accessible to all with basic laboratory facilities and mimics viability counts of *in vivo* models where *P. aeruginosa* is dominant and *S. aureus* is 2-3 log lower in abundance^29^. As such, it will be useful for fundamental studies of *P. aeruginosa* and *S. aureus* interactions. However, further investigation will reveal whether our findings apply within *ex vivo* host tissue models of infection, or within *in vivo* studies^41^. Examples of such *ex vivo* models include a model of primary ciliary dyskinesia, which incorporates *Haemophilus influenzae* in co-culture with diseased ciliated epithelial cells^42^, a model of *P. aeruginosa* utilising pig bronchioles^43^ or a dual species wound model of *P. aeruginosa* and *S. aureus* utilising immortalised keratinocytes as a substratum^44^.

In summary, we have described the creation and optimisation of a stable, co-culture biofilm model of *P. aeruginosa* and *S. aureus* and demonstrated that co-culture of these organisms can increase the rate of bacterial mutation, which could have important implications for studying bacterial evolution, adaptation and AMR within multispecies consortia.

## Acknowledgements

The authors would like to acknowledge the assistance of Steven Pullan of Public Health England for his assistance and expertise involving the nanopore sequencing of *P. aeruginosa* PAO1. The authors would also like to thank David Cleary (University of Southampton) for his valuable discussions surrounding analysis of the genomic data and whole genome comparisons and Qi Zheng (Texas A&M School of Public Health) for his advice surrounding the use of the rSalvador package and analysis of fluctuation analysis data. Short read genome sequencing was provided by MicrobesNG (http://www.microbesng.uk), which is supported by the BBSRC (grant number BB/L024209/1). In addition, the authors acknowledge the use of the IRIDIS High Performance Computing Facility, and associated support services at the University of Southampton, in the completion of this work. YM and AC made important contributions to the conception and design of the study and critiqued the output for important intellectual content.

This work was funded by a Biotechnology and Biological Sciences Research Council CASE Studentship award in partnership with Helperby Therapeutics, (BB/L016877/1). ARH was supported by NIH public health service grant AI083211. OS was supported by a Cystic Fibrosis Trust UK grant (Registered Charity No. for England and Wales 1079049) as part of a Strategic Research Centre titled “Pseudomonal infection in CF”.

## References

1. WHO. Antimicrobial resistance. Global Report on Surveillance. Bulletin of the World Health Organization 61, 383–94 (2014).

2. Koo, H., Allan, R. N., Howlin, R. P., Stoodley, P. & Hall-Stoodley, L. Targeting microbial biofilms: current and prospective therapeutic strategies. Nat. Rev. Microbiol. 15, 740 (2017).

3. Levin-Reisman, I. et al. Antibiotic tolerance facilitates the evolution of resistance. Science 355, 826–830 (2017).

4. Ahlgren, H. G. et al. Clinical outcomes associated with *Staphylococcus aureus* and *Pseudomonas aeruginosa* airway infections in adult cystic fibrosis patients. BMC Pulm. Med. 15, 67 (2015).

5. Serra, R. et al. Chronic wound infections: the role of *Pseudomonas aeruginosa* and *Staphylococcus aureus*. Expert Rev. Anti. Infect. Ther. 13, 605–613 (2015).

6. Phalak, P., Chen, J., Carlson, R. P. & Henson, M. A. Metabolic modeling of a chronic wound biofilm consortium predicts spatial partitioning of bacterial species. BMC Syst. Biol. 10, 90 (2016).

7. Mashburn, L. M., Jett, A. M., Akins, D. R. & Whiteley, M. *Staphylococcus aureus* serves as an iron source for *Pseudomonas aeruginosa* during in vivo coculture. J. Bacteriol. 187, 554–566 (2005).

8. Limoli, D. H. et al. Pseudomonas aeruginosa alginate overproduction promotes coexistence with *Staphylococcus aureus* in a model of cystic fibrosis respiratory infection. MBio 8, e00186–17 (2017).

9. Filkins, L. M. et al. Coculture of *Staphylococcus aureus* with *Pseudomonas aeruginosa* drives *S. aureus* towards fermentative metabolism and reduced viability in a cystic fibrosis model. J. Bacteriol. 197, 2252–2264 (2015).

10. DeLeon, S. et al. Synergistic interactions of *Pseudomonas aeruginosa* and *Staphylococcus aureus* in an *in vitro* wound model. Infect. Immun. 82, 4718–4728 (2014).

11. Duignan, C. M. Influence of low-dose nitric oxide on mono- and mixed-species biofilms formed by bacteria isolated from cystic fibrosis patients. (University of Southampton, 2017).

12. Hu, Y., Shamaei-Tousi, A., Liu, Y. & Coates, A. A new approach for the discovery of antibiotics by targeting non-multiplying bacteria: A novel topical antibiotic for Staphylococcal infections. PLoS One 5, e11818 (2010).

13. Howlin, R. P. et al. Antibiotic-loaded synthetic calcium sulfate beads for prevention of bacterial colonization and biofilm formation in periprosthetic infections. Antimicrob. Agents Chemother. 59, 111–120 (2015).

14. Foster, P. L. Methods for determining spontaneous mutation rates. Methods Enzymol. 409, 195–213 (2006).

15. Hall, B. M., Ma, C. X., Liang, P. & Singh, K. K. Fluctuation anaLysis calculator: A web tool for the determination of mutation rate using Luria-Delbück fluctuation analysis. Bioinformatics 25, 1564–1565 (2009).

16. Wick, R. R., Judd, L. M., Gorrie, C. L. & Holt, K. E. Unicycler: resolving bacterial genome assemblies from short and long sequencing reads. PLOS Comput. Biol. 13, e1005595 (2017).

17. Seemann, T. Prokka: rapid prokaryotic genome annotation. Bioinformatics 30, 2068–2069 (2014).

18. Overbeek, R. et al. The SEED and the Rapid Annotation of microbial genomes using Subsystems Technology (RAST). Nucleic Acids Res. 42, D206–14 (2014).

19. Aziz, R. K. et al. The RAST Server: Rapid Annotations using Subsystems Technology. BMC Genomics 9, 75 (2008).

20. Zheng, Q. New algorithms for Luria-Delbrück fluctuation analysis. Math. Biosci. 196, 198–214 (2005).

21. Heydorn, A. et al. Quantication of biofilm structures by the novel computer program. Image Process. 146, 2395–2407 (2000).

22. MacGregor-Fors, I. & Payton, M. E. Contrasting diversity values: statistical inferences based on overlapping confidence intervals. PLoS One 8, e56794 (2013).

23. Zheng, Q. Methods for comparing mutation rates using fluctuation assay data. Mutat. Res. - Fundam. Mol. Mech. Mutagen. 777, 20–22 (2015).

24. Hunter, R. C. et al. Ferrous Iron Is a Significant Component of Bioavailable Iron in Cystic Fibrosis Airways. MBio 4, e00557–13–e00557-13 (2013).

25. Tyrrell, J. & Callaghan, M. Iron acquisition in the cystic fibrosis lung and potential for novel therapeutic strategies. Microbiology 162, 191–205 (2016).

26. Noinaj, N., Guillier, M., Barnard, T. J. & Buchanan, S. K. TonB-dependent transporters: regulation, structure, and function. Annu. Rev. Microbiol. 64, 43–60 (2010).

27. Baldan, R. et al. Adaptation of *Pseudomonas aeruginosa* in cystic fibrosis airways influences virulence of *Staphylococcus aureus in vitro* and murine models of co-Infection. PLoS One 9, e89614 (2014).

28. Palmer, K. L., Mashburn, L. M., Singh, P. K. & Whiteley, M. Cystic fibrosis sputum supports growth and cues key aspects of *Pseudomonas aeruginosa* physiology. J. Bacteriol. 187, 5267–77 (2005).

29. Seth, A. K. et al. Comparative analysis of single-species and polybacterial wound biofilms using a quantitative, *in vivo*, rabbit ear model. PLoS One 7, e42897 (2012).

30. Miller, C. L. et al. Global transcriptome responses including small RNAs during mixed-species interactions with methicillin-resistant *Staphylococcus aureus* and *Pseudomonas aeruginosa*. Microbiologyopen 6, 1–22 (2017).

31. Nguyen, A. T. & Oglesby-Sherrouse, A. G. Spoils of war: iron at the crux of clinical and ecological fitness of *Pseudomonas aeruginosa*. BioMetals 28, 433–443 (2015).

32. Whittaker, J. W. Non-heme manganese catalase-the ‘other’ catalase. Arch. Biochem. Biophys. 525, 111–20 (2012).

33. Liu, M., Zhang, Y., Inouye, M. & Woychik, N. A. Bacterial addiction module toxin Doc inhibits translation elongation through its association with the 30S ribosomal subunit. Proc. Natl. Acad. Sci. U. S. A. 105, 5885–90 (2008).

34. Ryder, V. J., Chopra, I. & O’Neill, A. J. Increased mutability of *Staphylococci* in biofilms as a consequence of oxidative stress. PLoS One 7, e47695 (2012).

35. Driffield, K., Miller, K., Bostock, J. M., O’neill, A. J. & Chopra, I. Increased mutability of *Pseudomonas aeruginosa* in biofilms. J. Antimicrob. Chemother. 61, 1053–1056 (2008).

36. Tognon, M. et al. Co-evolution with *Staphylococcus aureus* leads to lipopolysaccharide alterations in *Pseudomonas aeruginosa*. ISME J. 11, 2233–2243 (2017).

37. Beaudoin, T. et al. Staphylococcus aureus interaction with *Pseudomonas aeruginosa* biofilm enhances tobramycin resistance. npj Biofilms Microbiomes 3, 25 (2017).

38. Radlinski, L. et al. Pseudomonas aeruginosa exoproducts determine antibiotic efficacy against *Staphylococcus aureus*. PLOS Biol. 15, e2003981 (2017).

39. Hubbard, A. T. M. et al. Mechanism of action of a membrane-active quinoline-based antimicrobial on natural and model bacterial membranes. Biochemistry 56, 1163–1174 (2017).

40. Orazi, G. & O’Toole, G. A. *Pseudomonas aeruginosa* alters *Staphylococcus aureus* sensitivity to vancomycin in a biofilm model of cystic fibrosis infection. MBio 8, e00873–17 (2017).

41. Lebeaux, D., Chauhan, A., Rendueles, O. & Beloin, C. From *in vitro* to *in vivo* models of bacterial biofilm-related infections. Pathog. (Basel, Switzerland) 2, 288–356 (2013).

42. Walker, W. T. et al. Primary ciliary dyskinesia ciliated airway cells show increased susceptibility to *Haemophilus influenzae* biofilm formation. Eur. Respir. J. 50, 1700612 (2017).

43. Harrison, F. & Diggle, S. P. An *ex vivo* lung model to study bronchioles infected with *Pseudomonas aeruginosa* biofilms. Microbiology 162, 1755–1760 (2016).

44. Alves, P. M. et al. Interaction between *Staphylococcus aureus* and *Pseudomonas aeruginosa* is beneficial for colonisation and pathogenicity in a mixed biofilm. Pathog. Dis. 76, (2018).

